# Evidence for strong mutation bias towards, and selection against, T/U content in SARS-CoV2: implications for attenuated vaccine design

**DOI:** 10.1101/2020.05.11.088112

**Authors:** Alan M. Rice, Atahualpa Castillo Morales, Alexander T. Ho, Christine Mordstein, Stefanie Mühlhausen, Samir Watson, Laura Cano, Bethan Young, Grzegorz Kudla, Laurence D. Hurst

## Abstract

Large-scale re-engineering of synonymous sites is a promising strategy to generate attenuated viruses for vaccines. Attenuation typically relies on de-optimisation of codon pairs and maximization of CpG dinculeotide frequencies. So as to formulate evolutionarily-informed attenuation strategies, that aim to force nucleotide usage against the estimated direction favoured by selection, here we examine available whole-genome sequences of SARS-CoV2 to infer patterns of mutation and selection on synonymous sites. Analysis of mutational profiles indicates a strong mutation bias towards T with concomitant selection against T. Accounting for dinucleotide effects reinforces this conclusion, observed TT content being a quarter of that expected under neutrality. A significantly different mutational profile at CDS sites that are not 4-fold degenerate is consistent with contemporaneous selection against T mutations more widely. Although selection against CpG dinucleotides is expected to drive synonymous site G+C content below mutational equilibrium, observed G+C content is slightly above equilibrium, possibly because of selection for higher expression. Consistent with gene-specific selection against CpG dinucleotides, we observe systematic differences of CpG content between SARS-CoV2 genes. We propose an evolutionarily informed gene-bespoke approach to attenuation that, unusually, seeks to increase usage of the already most common synonymous codons. Comparable analysis of H1N1 and Ebola finds that GC3 deviated from neutral equilibrium is not a universal feature, cautioning against generalization of results.

---

Understanding selective constraints on synonymous mutations (predominantly at codon third sites) provides the underlying rationale for the design of optimized transgenes (Fath, et al. 2011). For example, bacterial genes universally avoid strongly folded RNA structures at their 5’ ends (Gu, et al. 2010; Umu, et al. 2016), and this property has been used to optimize transgene expression (Kudla, et al. 2009; Goodman, et al. 2013; Boel, et al. 2016). Similarly, selection on synonymous sites causes coadaptation between codon usage and tRNA pools (Ikemura 1981; Higgs and Ran 2008), which can increase gene expression through a recently discovered mechanism that involves the recognition of slowly translated codons by a dedicated mRNA degradation pathway (Buschauer, et al. 2020). The effects of modification of synonymous site nucleotides are not modest: modulation of RNA structures, synonymous site GC content or codon-tRNA adaptation alters protein levels in prokaryotic and eukaryotic cells by orders of magnitude (Gustafsson, et al. 2004; Kudla, et al. 2009; Fath, et al. 2011; Bentele, et al. 2013; Mordstein, et al. 2020).

Understanding the mechanisms and role, if any, of selection on synonymous sites of viruses is similarly of applied importance as regards the design of attenuated versions for vaccines (e.g. Mueller, et al. 2010; Manokaran, et al. 2019; Cai, et al. 2020). In this instance, we aim to discern how selection acts on synonymous sites with a view to engineering the virus against the direction favoured by selection. In particular, viral attenuation can be achieved by alteration of synonymous sites as a means to modify the pattern of dinucleotides that bridge between successive codons (alias codon pair bias) while retaining the original protein (Karlin, et al. 1994; Rima and McFerran 1997; Coleman, et al. 2008). The same strategy has been extended to bacterial vaccines (Coleman, et al. 2011). Attenuation via modification of many synonymous sites has the notable advantage that any such virus employed as a vaccine will likely need many mutations to acquire wild-type fitness. Such a strategy is thus likely to be robust to virus/vaccine intra-host evolution. Indeed, given the mutation rate of SARS-COV2 (about 1 mutation every two weeks) this not a concern as regards intra-host adaptation by the virus. Synonymous codon manipulation has thus been proposed as a viable strategy for SARS-CoV2 attenuation and vaccine production (e.g. Kames, et al. 2020).

The codon pair bias attenuation effect has more recently been shown to be largely owing to increased CpG content (Tulloch, et al. 2014; Gaunt, et al. 2016). This is very likely to relate to the activity of the human Zinc Antiviral Protein (ZAP) as this targets transcripts with high CpG content (Takata, et al. 2017; Ficarelli, et al. 2020), although it is by no means the only antiviral peptide (Supplementary Table 1). As might be expected, ZAP is under positive selection owing to host-parasite coevolution (Kerns, et al. 2008). This suggests a simple attenuation strategy for SARS-CoV2, i.e. to increase CpG content (Kames, et al. 2020), this being consistent with the observed low CpG enrichment of the virus as sequenced in the wild (Xia 2020), also seen in cytoplasmic viruses more generally (Simmonds, et al. 2013). TpA is commonly considered alongside CpG not least because both are under-represented in native human transcripts (Simmonds, et al. 2013). Similarly, viruses lacking CpG also tend not to have TpA and engineering increased CpG and TpA attenuates viruses (Simmonds, et al. 2013). TpA depletion in SARS-CoV2 is weaker than CpG depletion (see below).

Given the possibility of selection against CpG and the possibility of being able to re-engineer synonymous sites, understanding forces operating on synonymous site composition, and nucleotide content more generally, is central to evolutionarily informed vaccine design, as well as to our understanding of the biology of SARS-CoV2. If there is selection against CpG residues, then we expect this to be evidenced as reduced G+C content at synonymous third sites, as mutation at third sites is least likely to have deleterious pleiotropic effects (e.g. via altering the amino acid specified). A codon with a C at position two selects for avoidance of G at the third site, while a codon starting with G selects for avoidance of C at the 5’ codon’s 3^rd^ site. In turn, we might expect that boosting third site G+C in a manner that increases CpG would be a viable attenuation strategy.

One means to test for such selection, or more generally fixation bias, is to identify a difference between predicted equilibrium G+C content under a neutral-mutation bias model and the values observed in the wild. To perform such a test one requires data on the relative rates of different classes of mutations (A->T, G->C etc) and from these rates per occurrence of the nucleotide calculate the equilibrium position i.e. the nucleotide content at which the rate of gain by mutation from other residues is equal to the rate of mutational loss. One can then compare observed and neutral equilibrium predicted values, with any discrepancy implicating a fixation bias.

Such methods have revealed commonplace deviations from null neutral expectations. For example, bacteria show a common GC->AT mutational bias (Hershberg and Petrov 2010), and hence a deviation from equilibrium in GC rich bacteria (Hildebrand, et al. 2010). Similarly, non-equilibrium TA nucleotide skews (Charneski, et al. 2011) have been identified. A recent large survey indicated that G+C deviating from neutral equilibria is common within eukaryotes (Long, et al. 2018). To derive this conclusion Lynch and colleagues extracted, from mutation accumulation (MA) experiments or parent-offspring sequencing, mutational profiles for numerous species and showed that the observed G+C content, even at codon third sites, was commonly higher than expected under a purely neutral model (Long, et al. 2018). The cause of this is unresolved, although GC biased gene conversion is one possible explanation (Long, et al. 2018). The possible importance of this process is supported by the widespread observation of a G+C-recombination correlation (Pessia, et al. 2012), implying higher GC in domains of more common double strand break repair. The case is by no means closed, however, as evidence for higher recombination in domains of manipulated high GC (Marsolier-Kergoat and Yeramian 2009; Kiktev, et al. 2018) and absence of GC:AT bias during gene conversion in organisms with a G+C recombination correlation (Liu, et al. 2018), both suggest the direction of the causative arrow may in some cases be reversed.

Rapid, accurate and common sequencing of epidemic and pandemic pathogens provide a rich source of data from which to derive the mutational profile (Hershberg and Petrov 2010; Hildebrand, et al. 2010; Charneski, et al. 2011). It is possible to ascribe both ancestral and derived states and hence infer the full mononucleotide mutational matrix (a 4 × 4, 12 parameter matrix of all possible mutations from one state to another) and, with enough mutations, the full dinucleotide matrix (a 16 × 16, 240 parameter matrix of all possible mutations from one dinucleotide to another).

As with parent-offspring sequencing and MA lines, we require that the mutations observed are an unbiased sample of the mutational profile (Long, et al. 2018). With very common sequencing (in all cases, short time periods between ancestor and progeny) we can ignore the possibility of multiple sequential hits at the same site (with the first hits going unsequenced) contaminating the mutational matrix. In principle the method can be misled by strong selection purging, in a non-random fashion, mutations prior their appearing in the population. However, if most selection is weak purifying selection there is then a lag between a deleterious mutation appearing (and being sequenced) and it being purged from a population. Declines in Ka/Ks as time to common ancestry increases in closely related bacteria strains (Rocha, et al. 2006) is consistent with such a model. In principle, even if there is strong selection on some mutations this too need not be problematic, so long as the profile of the mutations subject to strong selection is itself unbiased. Here then we apply this method to SARS-CoV2. To be cautious we consider only “high quality” mutations at four-fold degenerate sites to define the “cleanest” mutational matrix. We show that employment of mutations at all CDS sites makes a small but detectable difference, indicative of contemporaneous selection prior to sequencing on non-synonymous variants.

Under the assumption of selection against CpG (Xia 2020), we predict that the neutral mutational equilibrium GC content would be higher than the observed one. We find the opposite: the predicted neutral equilibrium (∼16.5%), owing to a strong GC->AT mutation bias, is lower than the observed one at synonymous sites (GC3∼28%, GC4∼20%). This is consistent with selection for raised G+C content but data suggest the more profound effect is selection against T content, as both T3 and TT we find to be far below their mutational equilibria. Regarding the rationale of any possible selection, we observe, even with a very underpowered test, a correlation between gene body nucleotide content and gene expression level that is suggestive of some expression mediated mode of selection. The direction of the effect is consistent with our prior observation that human intronless transgenes that are AT rich have poor expression potential (Mordstein, et al. 2020).

Unusually then, our data suggests that the most common third site residue (T) is also the one selected against. This result affects the strategy for attenuation. Classically we presume selection dominates over mutation bias and so to design highly expressed genes one selects for the common codons assuming them to be the most optimal. In this instance, a parsimonious interpretation of the data is that the mutation bias is so strong that selection has been unable to fully counter its effects, leaving a sub-optimal (selected against) nucleotide (T) the most commonplace. Given our results we thus propose the unusual strategy of increasing the usage of the already most highly used residue so as to degrade performance of the virus. Put differently, assuming selection acts against T and TT, a simple method to increase CpG content may not be the optimal strategy for viral attenuation. Given that prior evidence indicated that selection for reduced CpG content is particular to just immediate early genes (Lin, et al. 2020), we ask whether all SARS-CoV2 genes are equally subject to CpG bias. We find that they are not. Given this we propose a gene-bespoke approach sensitive to both CpG and putative selection on synonymous site T. Our results come with at least four caveats: that we cannot eliminate recently altered mutational biases, that we cannot address non-selective fixation biases, that sequencing errors may impact the mutational matrix and that we can only assume, but not demonstrate, that recent mutations observed at 4-fold degenerate sites are an unbiased reflection of the mutational process.

## METHODS

### Gene locations

We employed NC_045512 to specify the gene sequence to determine observed GC content, CpG content etc. However, following further annotation of genes (Kim, et al. 2020) we modified the gene locations to reflect those specified: https://github.com/hyeshik/sars-cov-2-transcriptome/blob/master/reference/SARS-CoV-2-annotations.gff. Specifically, to avoid a small codon overlap, we exclude the overlap hence employed annotation:

ORF7a protein 27394..27759 -> 27394..27753

ORF7b protein 27756..27887 -> 27762..27887

To consider ORF1a and ORF1b independently and to avoid overlap we employ:

ORF1a-> 266-13465

ORF1b -> 13471-21552

### Estimating flux rates from data

11,687 SARS-CoV-2 genome assemblies were downloaded from the GISAID (Shu and McCauley 2017) Initiative EpiCoV platform. Only assemblies flagged as “complete (>29,000 bp)”, “high coverage only”, and from a human isolate were downloaded. GISAID define “high quality” as entries with <1% Ns and <0.05% unique amino acid mutations (not seen in other sequences in database) and no insertion/deletion unless verified by submitter). Sequences were aligned with MAFFT 7.458 (Katoh and Standley 2013) to Wuhan-Hu-1 reference genome (EPI_ISL_402124). Sites in the first and last 200 bp of the alignment were masked to account for the fact that a higher degree of spurious variants tend to locate at the ends of the multiple sequence alignment. Variant sites were obtained from the MSA using the package SNP-sites (Page, et al. 2016) and whole genome nucleotide flux estimates were obtained by counting the frequency of each type of mutation with respect to the reference genome.

Isolates containing at least one coding sequence of length not divisible by three or containing a premature stop codon were excluded, removing 284 strains. CDSs were then translated using BioPython, re-aligned using MAFFT, and then reversed translated using TranslatorX (Abascal, et al. 2010). MSA of CDSs were concatenated and then, just as with the whole genome analysis, variant sites were obtained using SNP-sites and flux estimates were obtained by counting the frequency of each type of change with respect to the reference.

Additionally, H1N1 influenza A pdm09 sequences for strains collected between January 2009 and August 2010 that contained segments PB2, PB1, PA, HA, NP, NA, MP and NS were obtained from GISAID (Shu and McCauley 2017) for 4 segments: RNA polymerase subunit (PB2), hemagglutinin (HA), nucleoprotein (NP), and neuraminidase (NA). Sequences with length not divisible by three or containing a stop codon when translated were excluded. Remaining sequences were translated by BioPython and aligned to Mexican strain EPI_ISL_66702 using MAFFT, and reverse translated to nucleotides using TranslatorX (Abascal, et al. 2010).

Multiple sequence alignment of 1610 full Ebola virus (EBOV) genomes sampled between 17 March 2014 and 24 October 2015 in West Africa was downloaded from EbolaID database (Carneiro and Pereira 2016). The alignment includes the reference genome NC_002549.1. Genomes with a proportion of more than 10% missing sites were discarded. CDSs for each strain were obtained by extracting the coordinates from the reference genome on the alignment. In order to include in the analysis as the largest proportion of the gene ZEBOVgp4, the longest CDS (NP_066246.1) was used, and the shorter, overlapping proteins NP_066247.1 and NP_066248.1 were discarded. Just as in the case of H1N1, sequences with length not divisible by three or containing a stop codon when translated were excluded. Remaining sequences were translated aligned to the reference strain using MAFFT, and reverse translated to nucleotides using TranslatorX (Abascal, et al. 2010).

### Homoplasy screen

Sites can appear as having independently occurring mutations for at least two reasons: the extra mutation may be a sequencing error or it may be a true homoplasy (i.e. the same mutation at the same site occurring more than once independently) (van Dorp, et al. 2020). Sequencing errors need to be removed. In principle knowing how to handle true homoplasies in the construction of a mutational matrix is not as simple.

At first sight one might suggest that, as independent mutations, each occurrence of the mutation should be considered. The key question, however, is whether the mutational profile at these sites is representative of activity at other sites. If it isn’t, then their over inclusion will bias the matrix towards the profile of homoplasic sites away from that of the rest of the genome, which could itself cause a false signal of non-equilibrium status (i.e. where mutationally predicted and observed nucleotide compositions – largely at non-homoplasic sites-disagree). A *priori* by virtue of the fact that they are homoplasic we might suppose that mutational activity at these sites is not reflective of the mutational profile elsewhere in the genome and it is the equilibrium properties of other sites that we are interested in. One could then opt to filter out mutations at homoplasic sites considering them possibly unrepresentative. However, we don’t know they are unrepresentative and so their removal may be depleting the analysis of information. We also don’t know how many of the non-mutated sites had had the property of being homoplasic prior to current sequencing. An alternative, the middle way, is to include them but count all occurrences at any given site as one event, thereby employing the mutations but preventing such sites from overly skewing the matrix and further reducing the impact of possible (missed) sequencing errors.

We opt for the latter “middle way” approach but also check for resilience by removing such sites. Fortunately, as such sites are so rare (10 of 972 4-fold degenerate sites), removal of these sites makes no important difference to calculation of GC equilibrium content, nor to estimation of observed nucleotide content. We thus report the homoplasic-excluded results as minor asides. Phylogenetic tree of SARS-CoV-2 was downloaded from the COVID-19 Genomics UK Consortium website (https://www.cogconsortium.uk/, version of 24-04-2020). Subsequently, the MSA and the resulting tree were used to identify recurrent mutations (homoplasies) using HomoplasyFinder (Crispell, et al. 2019). All ambiguous sites in the alignment were set to ‘N’.

HomoplasyFinder identified 1740 putative homoplasies that were distributed over the SARS-CoV-2 genome. In order to remove spurious homoplastic sites that could arise due to sequencing error, these were filtered using a set of parameters and thresholds defined in (van Dorp, et al. 2020) to obtain a set of high confidence homoplasies. Briefly, for each homoplasy, the proportion of isolates with the homoplasy pnn where the nearest neighbouring isolate in the phylogeny also carried the homoplasy was computed and all homoplasies with pnn < 0.1 were excluded. Furthermore, we also excluded homoplasies that were shared in less than 0.1% of the isolates (>11 isolates). We also required that no isolate had an ambiguous base near the homoplasies (±5 bp). These filters reduced the number of homoplastic sites to 223.

### Estimating neutral equilibria

In principle one can estimate neutral GC equilibria knowing relative rates of GC->AT and AT->GC mutations alone e.g. (Long, et al. 2018). However, we take a fuller approach to estimate the equilibrium content of all nucleotides that also enables us to capture nucleotides skews (Charneski, et al. 2011). This has the advantage of treating all four bases as separate independent states, as is fitting for a single stranded virus unconstrained by base-pairing rules. Let us denote the frequency of G as G, the frequency of T, T etc. We shall write that the mutational frequency of G to T will be g2t etc, these being measured per occurrence of the starting base. The frequency of the nucleotides (N’) after some period will then be:

G’ = G (1-g2t-g2c-g2a) + A (a2g) + T (t2g) + C (c2g)

C’ = C (1-c2t-c2g-c2a) + A (a2c) + T (t2c) + G (g2c)

A’ = A (1-a2t-a2c-a2g) + G (g2a) + T (t2a) + C (c2a)

T’ = T (1-t2g-t2c-t2a) + A (a2t) + G (g2t) + C (c2t)

We then solve such that G’=G, T’=T etc. This thus resolves to:

G (g2t-g2c-g2a) = A (a2g) + T (t2g) + C (c2g)

C (c2t-c2g-c2a) = A (a2c) + T (t2c) + G (g2c)

A (a2t-a2c-a2g) = G (g2a) + T (t2a) + C (c2a)

T (t2g-t2c-t2a) = A (a2t) + G (g2t) + C (c2t)

Note that the left hand of each equation is the rate of loss given current abundance, while the right is the rate of gain given current abundances (i.e. we are solving for gain =loss). The 12 flux parameters (a2t, a2c etc) we derive from the mutational profile these being the number of observed changes per relevant occurrence of the nucleotide in the ancestral (pre mutated) sequence. We then solve these four simultaneous equations. Note that we replace any one arbitrarily chosen frequency by 1-sum of the other three (e.g. T= 1-A-C-G). These were solved in NumPy. Equilibrium solutions we denote with an asterisk (e.g. G*, GC3* etc). N4* implies nucleotide content of nucleotide N at 4-fold degenerate sites.

To assign bounds on the equilibrium estimates we perform a bootstrap test in which we resample with replacement M mutations from the set of M mutations. For each sampled vector we recalculate the predicted equilibria thereby assigning bounds. We report 95% bootstrap bounds from 100 re-samplings.

The same approach applies to the 16 × 16 dinucleotide matrix with 240 parameters.

### Estimating dinucleotide enrichment

For the dinucleotide NpM (e.g. CpG, GpC etc), we define gene body enrichment (E(NM)) as:

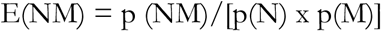

where p(NM) is the frequency of all dinucleotides within the gene that are NM and p(N) and p(M) are the frequencies of the mononucleotides within the same gene. We then consider site specific enrichment, i.e. sites 12, 23, or 31 defined by codon position, 31 being a third site and the codon first site of the following codon. Then at sites xy:

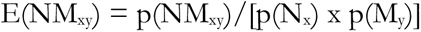

Where NM_xy_ is the relevant dinucleotide initiating with N at site x.

### Gene expression

We employ expression data specified by Kim et al. (2020). We used the highest read count for each subgenomic RNAs in Supplemental Table 3 of Kim et al. (2020) and compared log2 normalised read counts to gene G+C content, G+C at third sites, and CpG enrichment. Note that subgenomic RNA measures exclude ORF1a and ORF1b. ORF10 is excluded as no reads were identified. We used the shapiro.test function in R to test log2 transformed read counts for normality.

### Data compilation of vertebrate viruses

Vertebrate virus sequences were retrieved from the Virosaurus database (Virosaurus databases 2020_4.1, Release April 2020, file: Virosaurus90v 2020_4.1)(Gleizes, et al. 2020) [accessed 07 May 2020]. In this database, complete sequences were clustered at 90% to remove redundancy. Since in this database, herpesviridae and poxviridae are split in genes rather than full genomes, complete sequences for these viruses were retrieved from NCBI (refseq sequences). The same was also done for segmented viruses to allow calculation of sequence parameters per species. Genome classification was retrieved from ICTV (Master Species List #34 version 2, release May 2019)(Walker, et al. 2019). Annotation for replication compartments was assigned according to ICTV (Walker, et al. 2019) and ViralZone (Hulo, et al. 2011) CpG and UpA enrichment were calculated as above. For virus sequences obtained from the Virosaurus database, the mean was derived to obtain one value per species. For segmented viruses, segments were first concatenated before calculating sequence parameters. Species information and sequence parameters can be found in Supplementary Table 2.

### Genome sources

We acknowledge the sources of the genomes that we employed in Supplementary Table 3 (for SARS-CoV-2), Supplementary table 4 (for H1N1) and Supplementary table 5 (for Ebola).

### Comparing mutational matrixes

We sought to test whether the predicted equilibria solutions were different between the matrixes reflecting mutational profiles at 4-fold degenerate sites and all mutations at other sites (i.e. not 4 fold degenerate), as might be predicted were there contemporaneous selection against mutations that are non-synonymous. We partitioned all CDS mutations into those at 4-fold redundant sites (n=972) and all others (n=4672). Using these two datasets we calculated observed equilibrium frequencies for each nucleotide (4* for 4-folds and n4* for non 4-folds0, representing each as a vector of length four. We then determined the Euclidean distance between the two vectors. To test for significance, we compare the magnitude of this Euclidean distance to that expected by chance employing a non-parametric Monte Carlo simulation. To this end, we randomly extracted without replacement 972 mutations from the full set of mutations so as to create a subsample of pseudo ‘4-folds’. The remaining 4672 mutations we then considered a sample of pseudo ‘non 4-fold’ mutations. For each randomization, we assembled the corresponding mutational matrix, solved for equilibria and calculated the Euclidean distance between the resulting vectors of predicted equilibrium for the four nucleotides. We repeated this procedure 10,000 times to generate a null distribution of Euclidean distances that controls for sample sizes differences. Significance was given a P = n/m, where n is the number of simulations in which the Euclidean distance is as great or greater than observed in the real data and m is the number of simulations (i.e. 10,000). To check for robustness we considered an alternative distance metric, namely sum of modular differences (Euclidean distance considers square root of sum of squares of difference).

To consider each nucleotide individually, from the same Monte Carlo sampling, we calculated the difference between predicted equilibria at sampled pseudo ‘four-folds’ and pseudo ‘non four-folds’ for the 10,000 repeats. This generates four distributions, one for each nucleotide. For each nucleotide we calculate the mean (approximately zero) and standard deviation of these randomizations. The observed difference seen for each nucleotide between the equilibria predicted using mutations at four-fold sites (their predicted neutral equilibria) compared to that calculated using mutations at non four-fold site, may then be represented as a Z-score (Z=(observed – mean of simulations)/sd of simulations), Z >|1.96| indicating significant deviation.

## RESULTS

### SARS-COV2 mutations are heavily GC->AT biased

From the 11,687 genomes we can identify spontaneous mutations. From these we derive a mutational matrix and from this we solve for mutational equilibrium. From 972 mutations at four-fold degenerate third sites we find a heavily GC->AT biased mutational profile (Table 1). From this we deduce that equilibrium GC (termed GC*) should be 16.47% (95% bootstrap estimates 16.19-16.60). The corresponding number is 16.6% on removing 10 homoplasies. Specifically, we find: T4*=64.7%; A4* = 18.93; C4* =12.28%; G4* = 4.19%.). The striking bias towards T has been recently commented on and considered to be consistent with APOBEC editing (Simmonds 2020).

**Table 1.** The 4 × 4 mutational matrix for 972 mutations at four-fold synonymous sites (in bold) and from 5644 mutations observed anywhere in codons (not bold). Rates are defined as the number of observed changes per incidence of the nucleotide in the reference genome at four-fold third sites (bold) or in codons. Note that because of different normalizations, the two sets of numbers are not directly comparable in absolute terms.

| Reference allele | Derived allele |  |  |  |  |
| --- | --- | --- | --- | --- | --- |
|  |  | A | T | C | G |
|  | A | - | **0.05041**<br>0.02043 | **0.01707**<br>0.01504 | **0.09350**<br>0.07771 |
|  | T | **0.02269**<br>0.01668 | - | **0.09398**<br>0.07340 | **0.01343**<br>0.01126 |
|  | C | **0.05842**<br>0.03265 | **0.45704**<br>0.35280 | - | **0.01375**<br>0.00970 |
|  | G | **0.20652**<br>0.10434 | **0.43841**<br>0.15373 | **0.02536**<br>0.01867 | - |

Cognisant that there might be dinucleotide based mutation biases we extend the mononucleotide matrix to a 16 × 16 dinucleotide matrix with 240 parameter estimates derived across the coding sequences (Fig 1, Supplementary table 6). With 11073 dinucleotide switches this represents an average of 46.1 mutations per parameter estimate which is liable to be noisy and potentially weakly influence by selection on non-synonymous mutations. With this we determine equilibrium content for all dinucleotides and in turn all nucleotides (A*=0.187, C*=0.112, T*=0.620, G*=0.0812). We thus estimate from this GC* of 19.3% (95% bootstraps 0.193-195) more or less in line with mononucleotide calculations.

**Figure 1.**
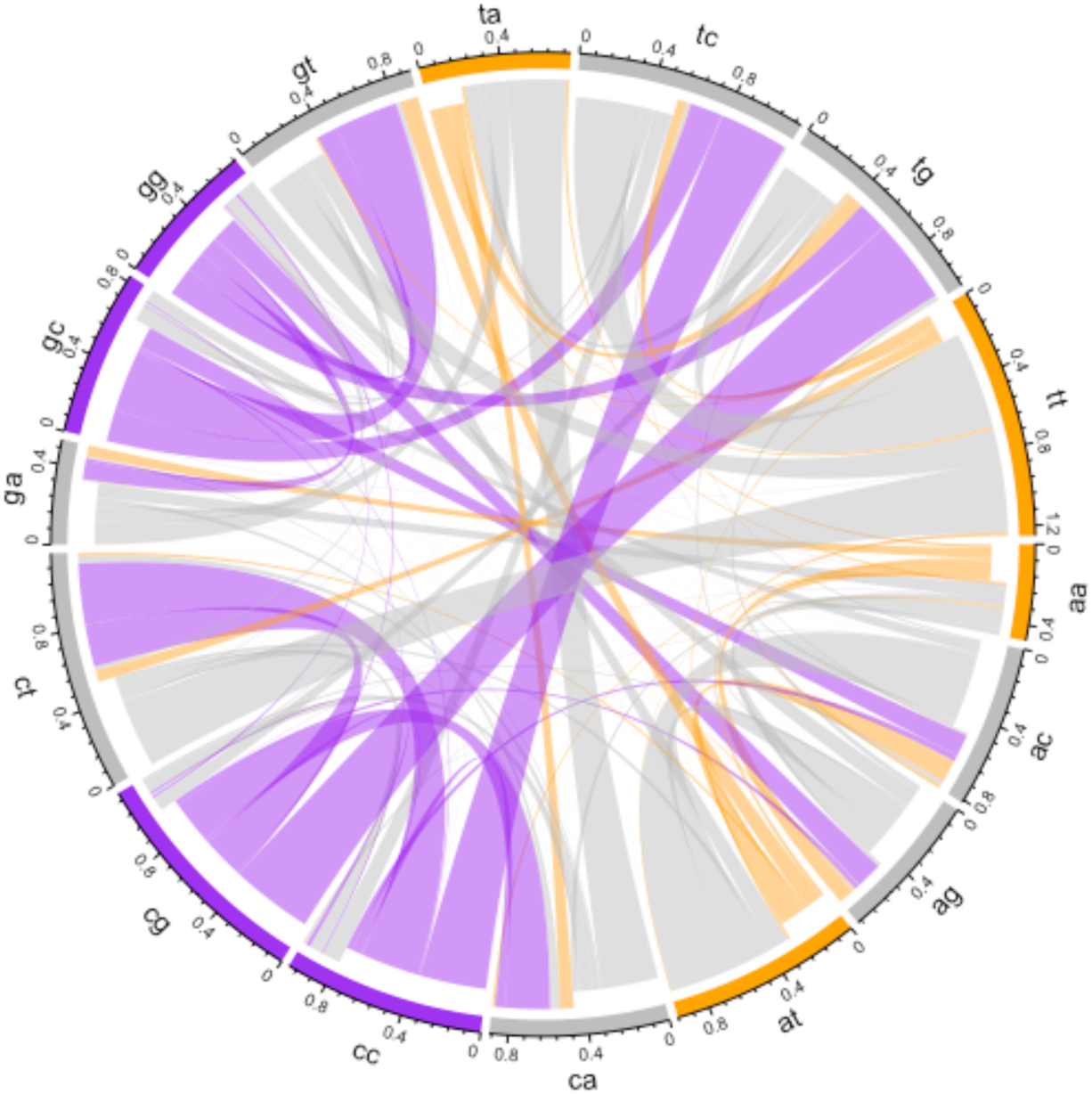
Chord diagram displaying the rate of flux from one dinucleotide to another in the coding sequence of SARS-CoV-2. For each node, the direction of flux is indicated by the indentation of the connecting links: the outer most layer represents flux into the node and the inner layer represents flux out. The frequency of the flux exchange is represented by the width of any given link where it meets the outer axis. Dinucleotide nodes are coloured according to their GC-content. Hence, it is evident that there is high flux away from GC-rich dinucleotides whereas AT-rich dinucleotides are largely conserved.

### Evidence for selection acting to counter a large mutation bias towards T

If selection favours reduced G+C content owing to selection for reduced CpG content, we expect that the observed GC3 should be lower than that predicted under neutrality (16.5%). We find the opposite to be true, observed GC3 being 28% (GC3 at 4-fold sites = 20.2%). All numbers are beyond 95% bootstrap bounds of the predicted equilibrium frequency derived from analysis of mononucleotide profiles at 4-fold degenerate sites (bounds: 16.19-16.60). More specifically, at four fold synonymous sites, observed T4 (50.8%) is less than predicted under neutral equilibrium T4* (64.6%) while all other bases are higher than expected (A4 = 28.95%, A4* = 18.93; C4 = 13.70%, C4* =12.28%; G4 = 6.50%, G4* = 4.19%.). A parsimonious explanation is that the sizeable mutation bias towards T generates deleterious mutations, non-optimal even at synonymous sites, and selection therefore favours reduced T content. However, increasing C or G potentially comes at a cost of increased CpG, so the base most in excess of its equilibrium is A.

GC of coding sequence is even more removed from the neutral equilibrium at 38%. This suggests selection in favour of non-synonymous mutations that increase G+C content. Examination of non-equilibrium status by dinucleotide content supports this. It shows one striking effect, namely that TT’s predicted equilibrium frequency greatly exceeds what is observed (Figure 2). More generally, T content whether derived from mononucleotides at 4-fold third sites (predicted 64.6%) or mononucleotides across the genes (predicted 59%) or from dinucleotides (62%) is greatly in excess of T content this being 32% for the complete viral sequence. The mutational matrix, whether through mono or dinucleotide analysis, predicts a great enrichment of T which we infer is being opposed by selection at third sites and in gene bodies (unweighted gene body means: T1% = 25.7%, T2%=36.3%, T3%=41%). We notice that CpG content is above that expected under neutrality (Figure 2). However, this we suggest is not so much evidence against selection towards high CpG so much as selection against TT which by necessity increases the observed relative frequency of CpG and most other dinucleotides as frequencies must sum to one.

**Figure 2.**
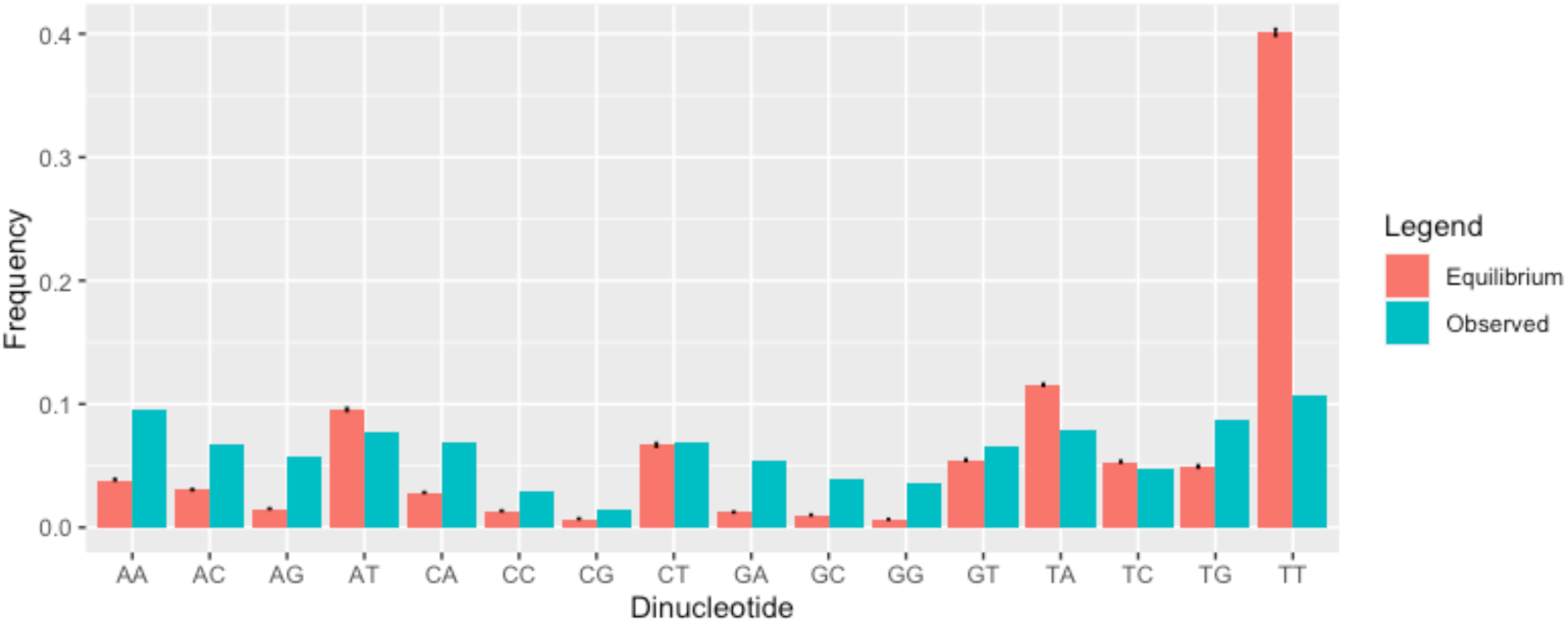
Comparison of dinucleotide content across SARS-CoV2 compared with neutral expectations. Error bars represent bootstrapped 95% upper and lower confidence bounds.

### Evidence for contemporaneous selection against T at non-four-fold redundant sites

Possibly consistent with a role for selection, using 5644 mutations that occur anywhere in the coding sequence (Table 1), we observe that the G->T flux at 4-fold degenerate sites is much greater than that observed throughout the sequence. Assuming the flux rate at 4-fold degenerate sites is more indicative of the true mutational flux, this is consistent with non-synonymous T mutations being under strong enough selection to be eliminated prior to sequencing. The predicted bias from this matrix is thus slightly more GC rich than that determined from 4-fold redundant sites (GC* at 20.65%: 95% bootstrap estimates 20.60-20.78, 20.65% also after excluding homoplasies).

This difference is perhaps a surprising result as classically when considering very recent mutations we presume an absence of bias in the profile of mutations. The skew would be consistent with contemporary selection on non-4-fold sites opposing mutations towards T, consistent also with the different between T4*, T4 and T content overall. To ask whether the difference between the two equilibria solutions is significantly different, we developed a non-parametric Monte Carlo simulation (Methods). We find that the Euclidean distances from the random sampling are the same as, or greater than, the Euclidean distance between four-folds and non-four fold sites in just 390/10,000 cases (hence P = 0.039) (repeating using an alternative distance metric, sum of modular difference between equilibria, make no meaningful difference, P=0.4). Thus, not only do we detect deviation away from the predicted neutral equilibrium (at 4-fold sites, third sites generally and through the gene body), we even can detect a signal consistent with selection on SARS-CoV2 that skews the mutational matrix prior to sequencing.

To clarify that it was selection against T, we considered each nucleotide individually (see Methods). Such analysis indeed provides evidence for significant counter selection of T at non-4-fold sites (Z = −2.01). Commensurably, predicted G equilibrium content derived from mutations at non-4-fold sites is higher than that derived from mutations at 4-fold degenerate sites (Z = 4.92), while A and C content are less affected (Z for A = 0.47, Z for C = −0.50). Contemporaneous selection opposing G->T mutations at non-4-fold sites is a parsimonious explanation. As N->T mutations at codons NAA, NGA and NAG will generate stop codons (where N can be A, C or G), part of the selection at non-our-fold sites against mutations generating T may be selection against nonsense mutations. The nine codons should be at a frequency of 9/61 =14.75% under unbiased nucleotide content but are at 17.05% with AAA (3.76%) being the second most common codon after GTT (3.9%).

### Significant heterogeneity in the degree of CpG avoidance between genes

While selection against T or TT provides a viable model for GC3>GC3*, might there be other explanations that would be consistent with selection against CpG, to avoid ZAP, but in favour of G+C? One possibility is that we may be witnessing between-gene heterogeneity (Digard, et al. 2020). Imagine that some genes are indeed under selection for low CpG and hence for low GC3, but others are not under selection for low CpG and thus are more free to have selection favouring higher GC3 (for unspecified reasons, but possibly to enable efficient expression (Mordstein, et al. 2020)). When then considered *en masse* we see both selection for CpG and more raised GC3. Recent reports suggest that not all genes are equally subject to selection for low CpG to avoid ZAP, with only “immediate early” genes under such selection (Lin, et al. 2020).

Were this the explanation, or part thereof, we would predict that CpG enrichment would be heterogeneous between genes (see also Digard, et al. 2020) and that those with relatively high CpG enrichment will also be those genes contributing to raised GC3 (i.e. a positive correlation between CpG enrichment and GC3). Note that while CpG counts are likely to be necessarily higher as GC3 goes up, CpG enrichment is normalised to underlying GC content and so CpG enrichment and high GC3 are not logically coupled (e.g. if at the limit 50% of residues are C and 50% G, so long as CpG usage is random, CpG = 0.5*0.5, CpG enrichment will not be seen).

To assay this, we calculated CpG enrichment at codon sites 12, 23 and 31, these providing three measures of CpG enrichment for each gene. We can then perform a Kruskal-Wallis test for heterogeneity. Even with such limited data, we find that the three measures for the same gene are more similar than expected by chance (KW, P=0.019, df=11: mean E(CG) =0.61 +/- 0.4 sd; Fig 3a). This implies that at all sites CpG is avoided or preferred to the same degree within any given gene. We see however only marginal evidence that genes released from CpG constraint are those with higher GC3 (CpG enrichment v GC3, rho =0.41, P=0.19, Spearman’s test, Fig 3b). Thus, while there is evidence for differential CpG usage between genes, we don’t find that this predicts GC3, although trends are in the expected direction and the tests underpowered.

**Figure 3.**
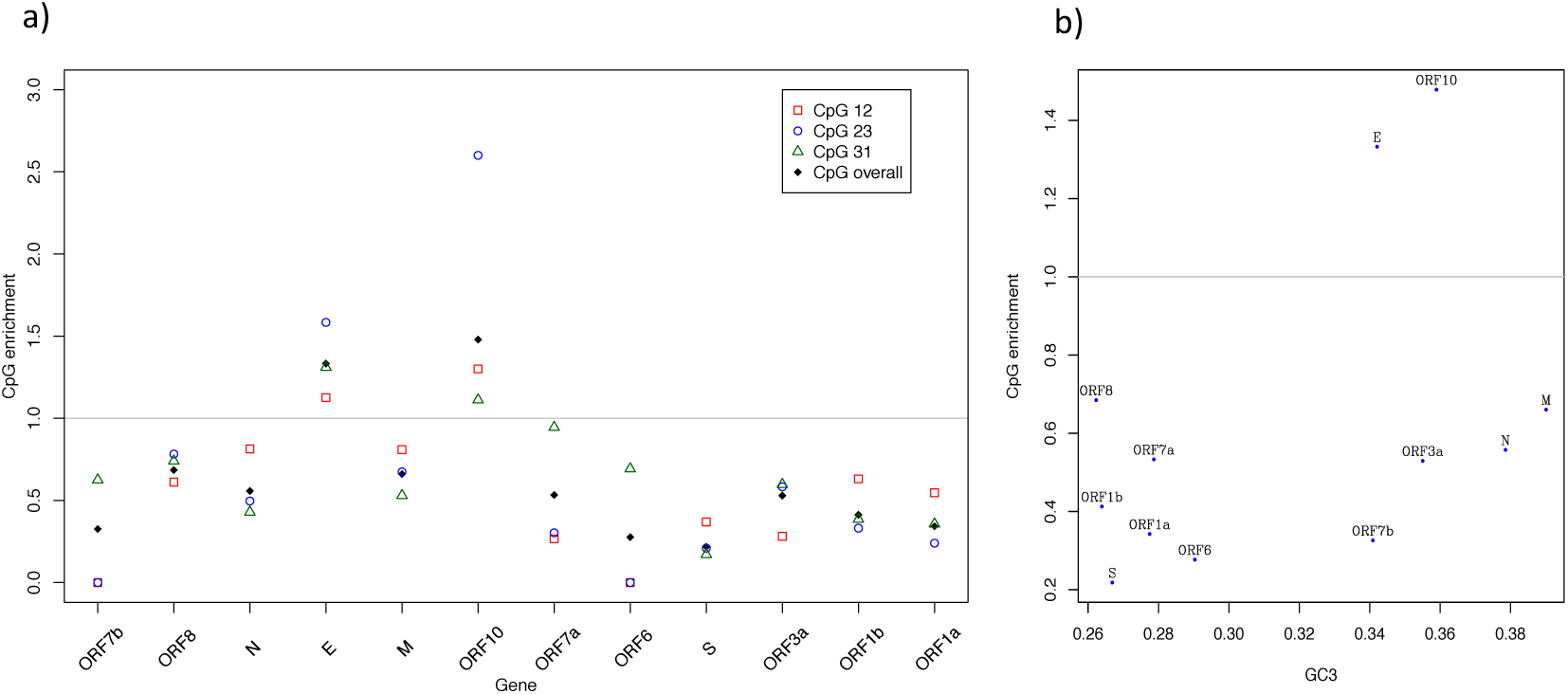
**a)** CpG enrichment across the genes of SARS-CoV2. Grey line = no enrichment **b)** relationship between CpG enrichment and GC3

More generally we can ask whether gene body G+C content behaves the same as gene body CpG content with each gene having its own characteristic profile. We assay this by considering GC1, GC2 and GC3 in a manner as above. We find no evidence that genes are more similar in these three measures than expected by chance (KW P=0.49, df=11: Fig 4). Similarly, we see no correlation between GC3 and GC12 although the trend is positive (rho = 0.15, P=0.63, Spearman’s rank). However, we do observe some regularities. First, GC3 is consistently lower than GC12 (Wilcoxon signed-rank test, P=0.007), the mean GC3 being 28%, while that of GC12 is 40%, consistent with selection on amino acid content.

**Figure 4.**
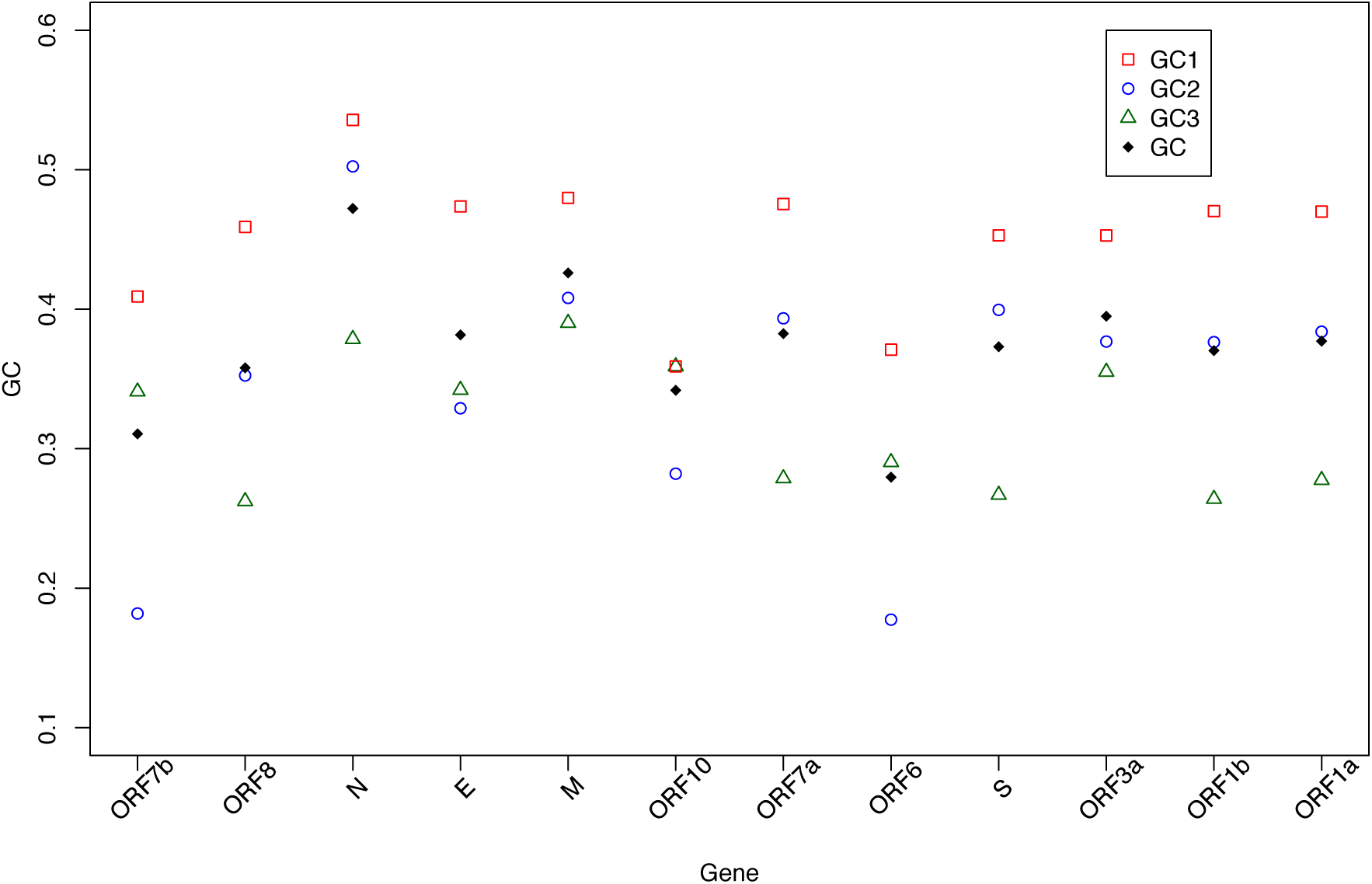
GC content across genes of SARS-CoV2 at codon sites 1, 2 3 and averaged across the gene

The most striking feature of third site nucleotide usage is that all genes have a preponderance of T (Figure 5). As noted above, this we can attribute only in some part to mutation as the predicted levels while in the rank order as observed (T>A>C>G) are highly deviant from null. Specifically, the predicted numbers are 0.65>0.19>0.12>0.04 while the observed are 0.44>0.28>0.16>0.13. Approximately the same predicted equilibrium values are seen employing all mutations (0.59>0.20>0.12>0.08). Selection against T seems strong, despite this being the most common nucleotide, as it is heavily reduced from its predicted equilibrium content.

**Figure 5.**
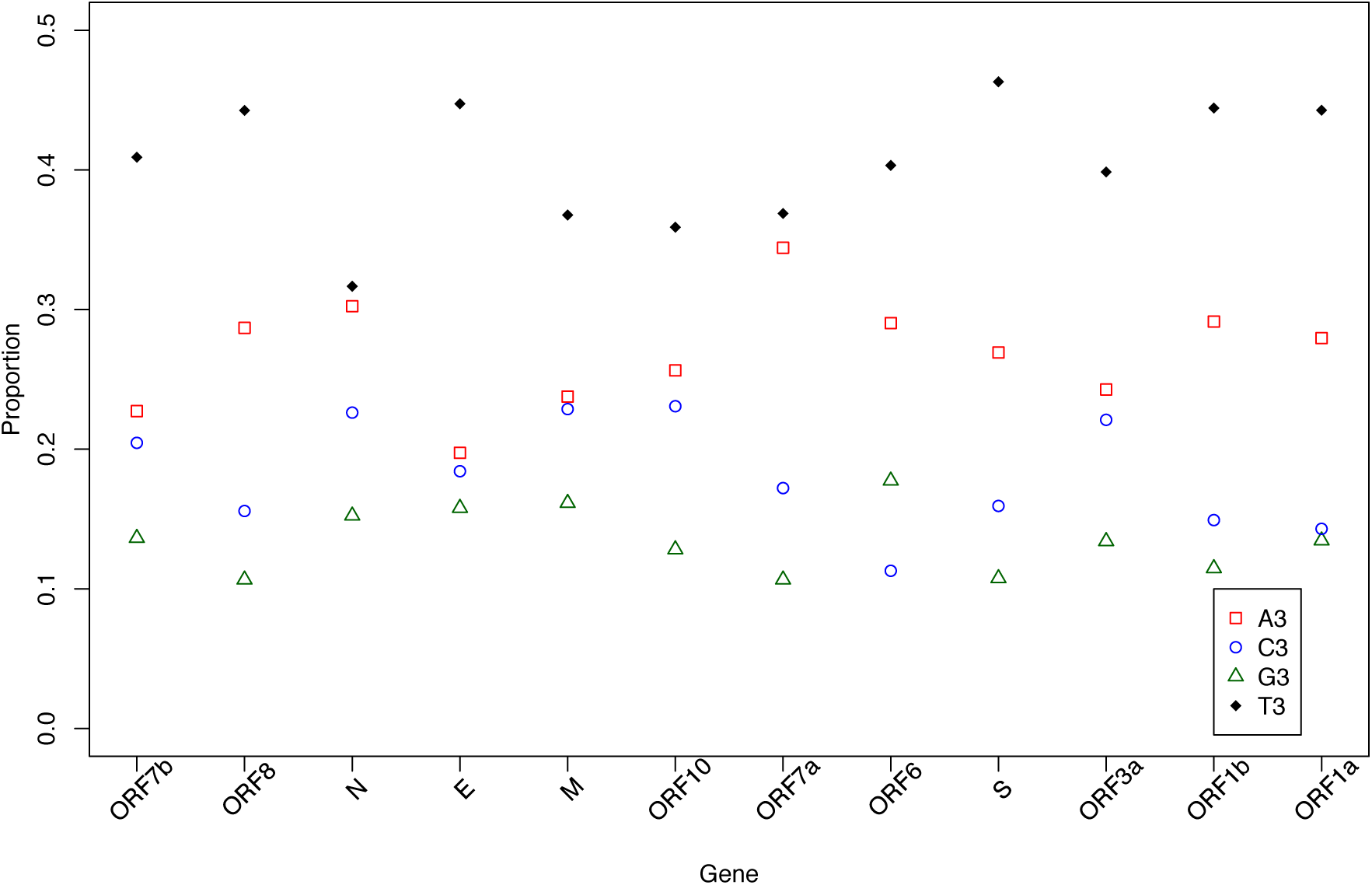
Base composition at codon third sites across genes of SARS-CoV2.

### Genes avoiding CpG also avoid TpA

Prior analysis suggests that viruses lacking CpG also tend not to have TpA and that engineering increased CpG and TpA attenuates viruses, possibly because both are under-represented in human transcripts (Simmonds, et al. 2013). We also observe that TpA enrichment and CpG enrichment tend to positively correlate across viruses (N=1344, rho=0.17, P=2.34 × 10^-10^: data in Supplementary Table 2). More particularly, this is seen only (or more profoundly) within the class of viruses, like SARS-CoV2, that cytoplasmically replicate (cytoplasmic: rho=0.12, P=0.0016, N=654; nuclear: rho=0.069, P=0.07, N=690). To understanding whether increasing CpG and TpA might be a useful attenuation strategy, we ask whether TpA is also avoided in genes of SARS-CoV2 and whether it is avoided in the same genes that avoid CpG. We consider not just the CpG enrichment predicting TpA enrichment but also, to control for mononucleotide effects, the two other symmetric nucleotide pairings (ApT and GpC).

On the average TpA is, like CpG, avoided although not to the same extent as CpG (mean TpA enrichment =0.83 +/- 0.2 sd) (Figure 6b). TpA also shows between gene heterogeneity (KW test P=0.04). We find that exclusively for CpG enrichment and TpA enrichment do we see a correlation between genes (Table 2, Fig 6a). ApT is also avoided (mean enrichment = 0.83 +/- 0.14 sd), but there is no evidence for within gene homogeneity (KT test P=0.14) (Fig 7). By contrast, there is no evidence for GpC avoidance: mean GpC enrichment =1.13, +/- 0.34 sd, Fig 7) and genes do not show gene-specific GpC enrichment (KW, P=0.11, comparing GpC enrichment at sites 12, 23 and 31). We conclude that if CpG enrichment is a viable strategy to attenuate a gene, increasing TpA may also (although for reasons unknown).

**Figure 6.**
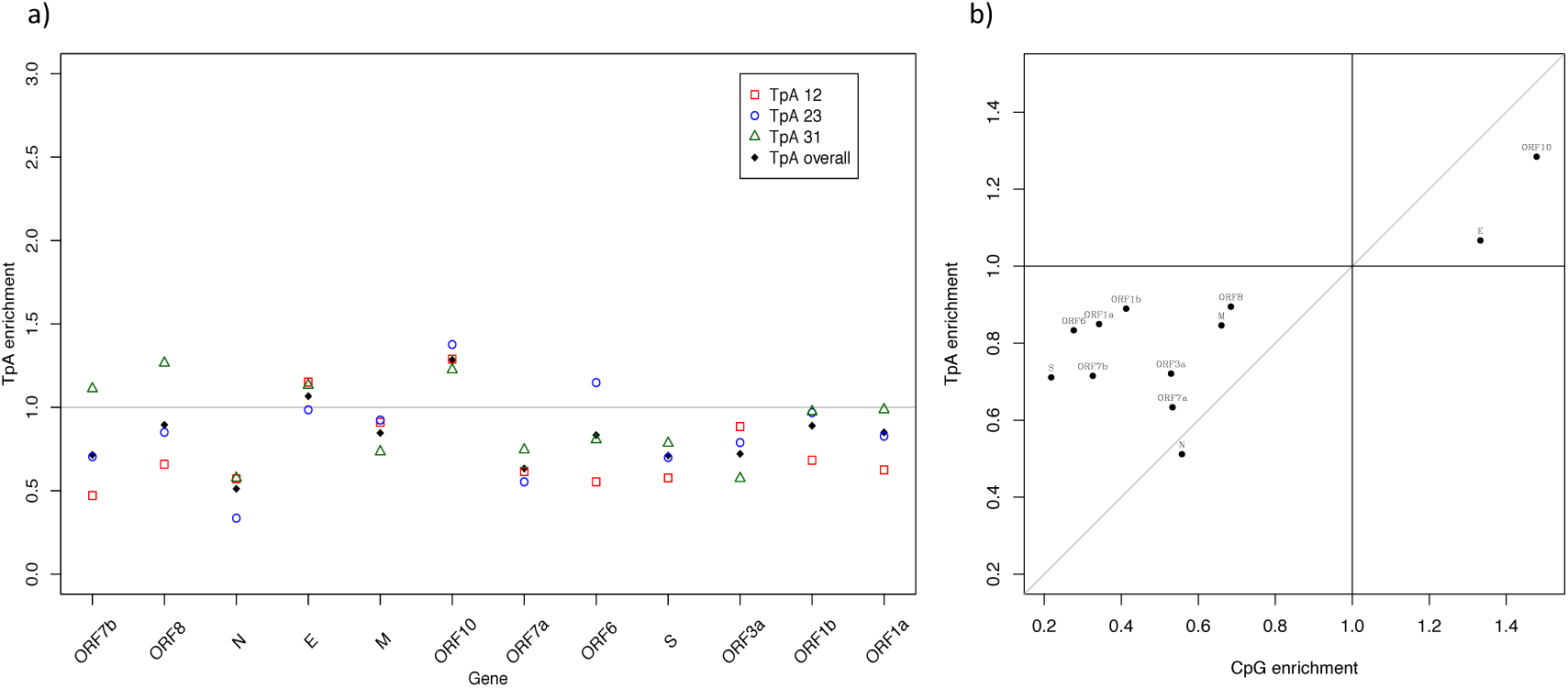
**a)** TpA enrichment across genes of SARS-CoV2 b) correlation with CpG enrichment. Grey line is the line of slope 1 through the origin.

**Figure 7.**
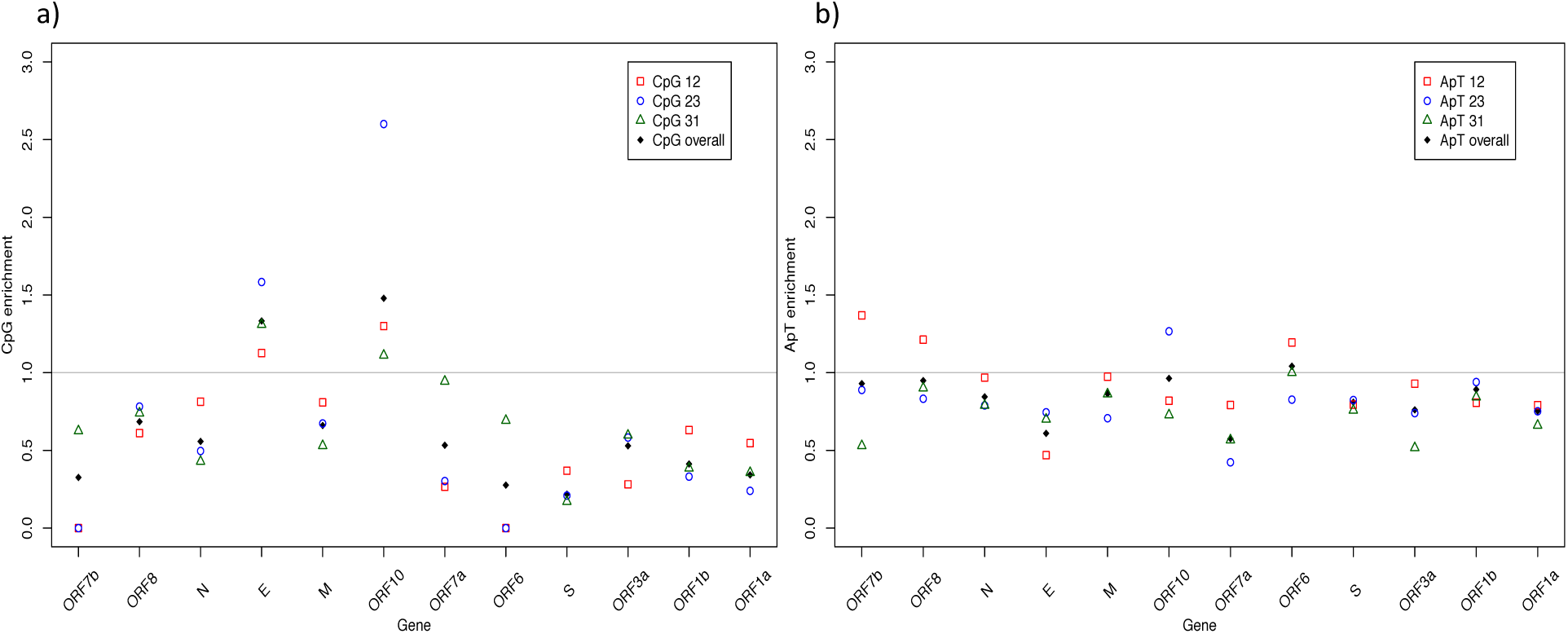
a) GpC and b) ApT enrichment across the genes of SARS-CoV2.

**Table 2.**
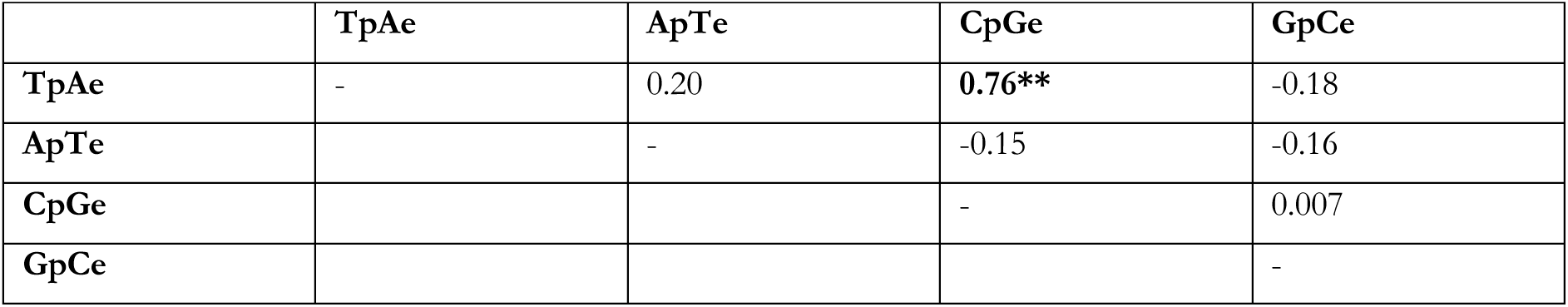
Between-gene correlations in dinucleotide enrichment scores (Pearson product moment correlation r values). Significant correlations in bold: ** = P <0.005.

### Evidence for T content predicting expression level

The results above are consistent with a model in which CpG content is under selection in some genes to be reduced, while GC3 content is above the level expected under neutrality, in no small part because the T mutation bias is so extreme that equilibrium T content (especially TT content) would render the virus much less fit. There are several possible mechanistic explanations for the GC3 > GC3* effect. With our recent evidence that intronless low GC genes barely express in human cell lines (Mordstein, et al. 2020), selection for raised GC3 (reduced T3) to enable more effective gene expression is a strong contender. In this context, while we do not see a GC3 expression correlation (r=0.09, P=0.82), we do observe a GC expression correlation (r=0.79, p=0.01 and figure 8). Breaking this down by nucleotide we see that this is owing to a negative correlation with T content and a positive correlation with both C and G content (A freq: r= 0.33, P=0.83; C freq: r=0.64, P=0.06; G freq r=0.81, P=0.009; T freq r=-0.88, P=0.0017). Why this is will require considerable experimental manipulation of sequences to understand. It is notable that we observe such an effect with such an underpowered test.

**Figure 8.**
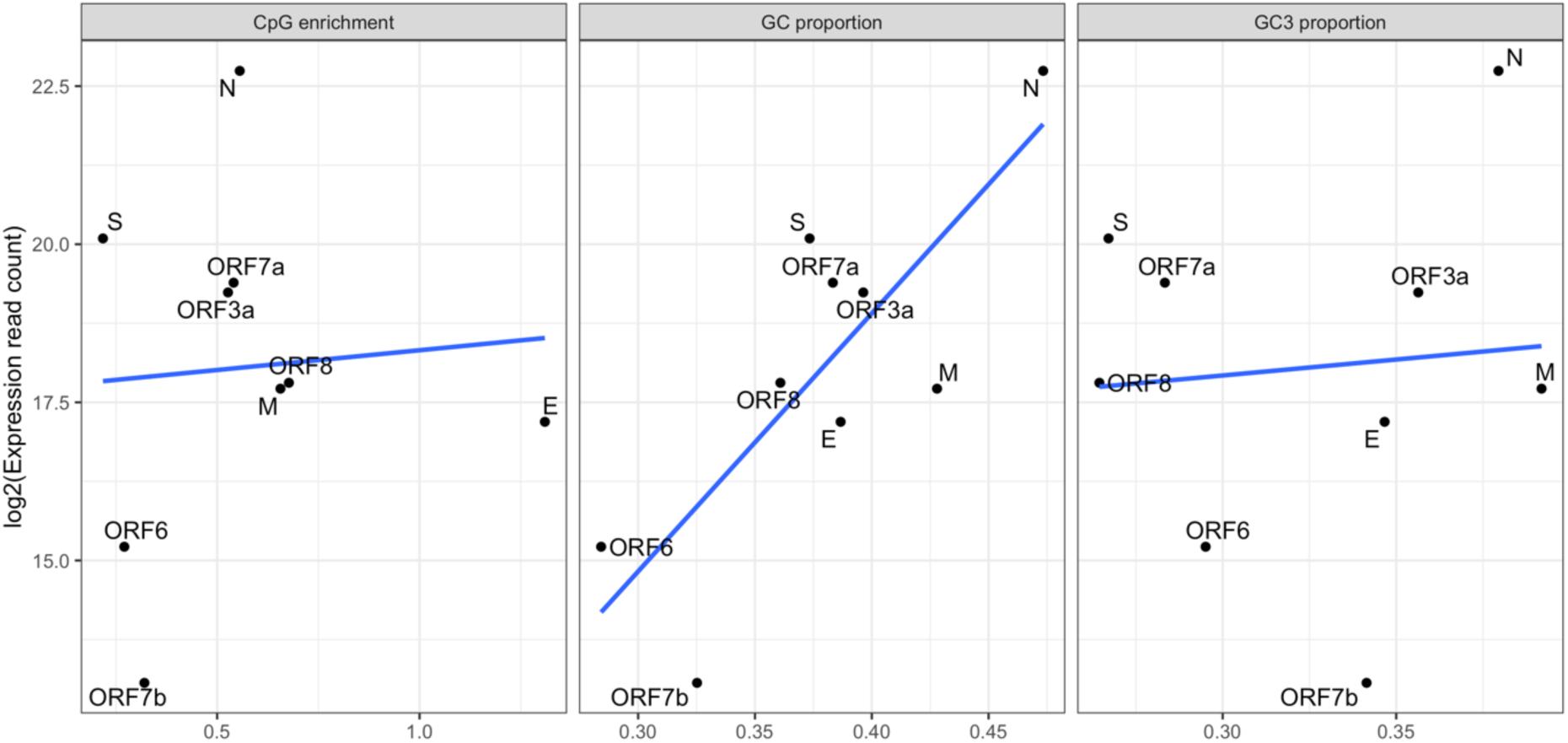
Correlation between expression level and CpG enrichment, GC content and GC3

A more broad-brush approach is to consider viral sequences more generally. As part of the mechanism by which GC enrichment boosts expression is thought to be intranuclear (e.g nuclear export) (Mordstein, et al. 2020), if selection is operating on gene expression of viruses, we might predict that nuclear viruses might have a higher GC content than cytoplasmic viruses. Using mean GC of all viruses within a taxonomic grouping we observe this to be the case (Mann Whitney U test P=1.6 10^-20^, Fig 9). CpG enrichment and TpA enrichment is similarly lower in cytoplasmic viruses (Fig 9). This is a very arms-length result and requires due caution in its interpretation (it could just as well be evidence of different mutational biases). Nonetheless, within the context of our prior result we suggest that this merits further scrutiny.

**Figure 9.**
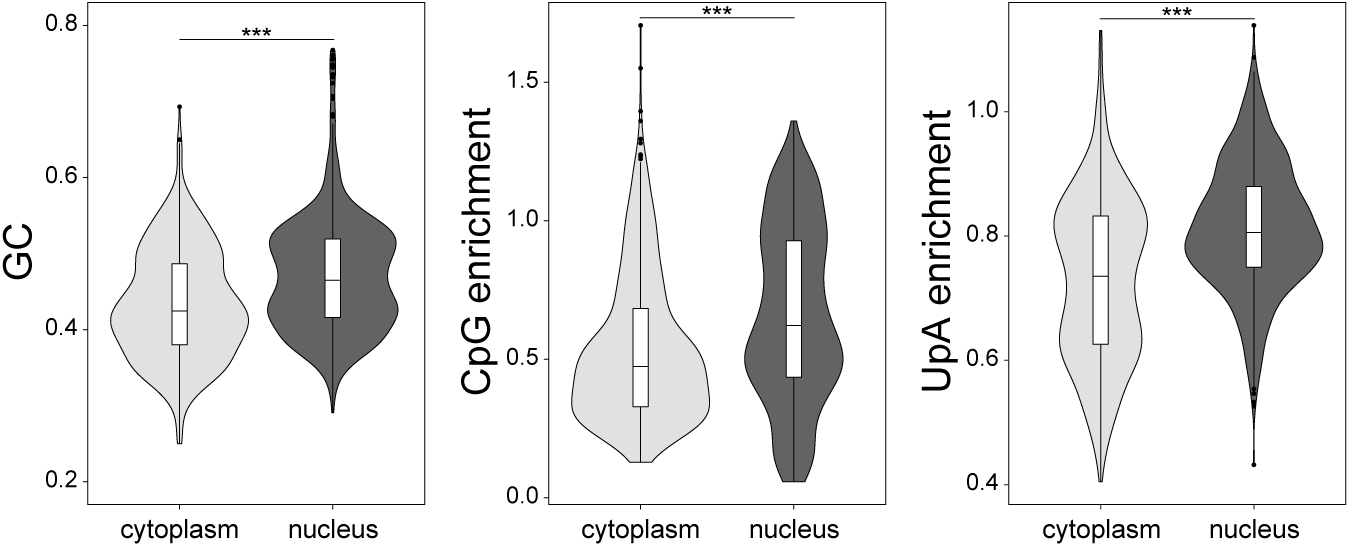
GC content of cytoplasmic and nuclear viruses. Cytoplasmic viruses have significantly lower values for all three measures (MWU test: GC: p = 1.63e-20, CpG enrichment p = 2.57e-15, [U/T]pA enrichment: p = 3.2e-36)

### Designing the optimally attenuated SARS-COV2

With the above evidence for selection for G+C at third sites and for heterogeneity between genes in enrichment of CpG and TpA, we suggest that simply increasing CpG by manipulation of synonymous sites need not be the optimal strategy. It may enable recognition by ZAP, but may also favour increased fitness by increasing G+C/reducing T (although we see no expression correlation employing just 4-fold degenerate sites although a negative T4 expression correlation is observed, but not significant: r=-0.38, p=0.32).

As not all genes are under selection for reduced CpG/TpA, reducing their G+C content by increasing T content seems a relatively safe and robust strategy. We thus suggest to classify genes according to the CpG enrichment: >1, 0.5-1 and <0.5. For those in the first category, likely not affected by ZAP (E and ORF10, (see also Digard, et al. 2020)) we suggest deceasing their synonymous G+C by increasing where possible T content and forcing them closer to their mutational equilibrium. For those with especially low CpG enrichment and most likely strong targets of ZAP (ORF1a, ORF1b, ORF6, ORF7b and S) we suggest, raising their CpG, even at the cost of increased G+C. Where possible TpA should also be increased. For the remainder we suggest maximizing CpG content while holding GC3 content static or decreasing if possible. However, with the possibility of synonymous sites also being parts of key motifs, e.g. for RNA binding proteins (Savisaar and Hurst 2017), a simplistic strategy, even if gene-tailored, may have deleterious undesirable side consequences.

### GC3*>GC3 is not a general property of viruses

We observed that GC content at third sites was both higher than expected given selection against CpG and higher than expected given the underlying mutational profile. Is a deviation from mutational equilibrium a general property of human viruses? Were this so, this too could have implications for engineering of attenuated forms. To address this, we consider other viruses with rich sequencing from epidemics.

For H1N1 using the same mode of analysis we observe both a less extreme GC->AT mutation bias (Table 3) and an observed GC3 content very close to that predicted. From analysis of 3rd sites the predicted value is GC3*=41.8% (bootstrap 95% intervals 41.45-42.04), from all sites the prediction is GC*=42.8% (bootstrap 95% 42.56-42.96). The observed GC3 is 41.8%, within the bounds of the prediction based on 3^rd^ site mutations. For Ebola (Table 4) we find observed GC3 is all but identical to predicted (observed GC3=46.4%, expected=46.7%). We conclude that analysis of SARS-CoV2 and its non-mutational equilibrium status at synonymous sites does not necessarily hold lessons for other viruses. In contrast to others (Kames, et al. 2020), we suggest caution in generalizing vaccine strategies.

**Table 3.** The 4 × 4 mutational matrix for 1522 mutations at synonymous sites (in bold) and from 2571 mutations observed anywhere in codons (not bold) for H1N1. Rates are defined as the number of observed changes per incidence of the nucleotide in the reference genome at 3^rd^ sites (bold) or in codons.

|  | Derived allele |  |  |  |  |
| --- | --- | --- | --- | --- | --- |
| Reference allele |  | A | T | C | G |
|  | A | - | **0.0871**<br>0.04597 | **0.065**<br>0.0451 | **0.4291**<br>0.25542 |
|  | T | **0.0803**<br>0.05143 | - | **0.4945**<br>0.24889 | **0.0529**<br>0.03429 |
|  | C | **0.1691**<br>0.11426 | **0.5699**<br>0.30675 | - | **0.0251**<br>0.02607 |
|  | G | **0.6089**<br>0.32052 | **0.0948**<br>0.05027 | **0.0323**<br>0.0207 | - |

**Table 4.** The 4 × 4 mutational matrix for 1682 mutations at synonymous sites (in bold) and from 3523 mutations observed anywhere in codons (not bold) for Ebola. Rates are defined as the number of observed changes per incidence of the nucleotide in the reference genome at 3^rd^ sites (bold) or in codons.

|  | Derived allele |  |  |  |  |
| --- | --- | --- | --- | --- | --- |
| Reference allele |  | A | T | C | G |
|  | A | - | **0.0739**<br>0.05077 | **0.0964**<br>0.06722 | **0.2123**<br>0.14803 |
|  | T | **0.0594**<br>0.05152 | - | **0.2145**<br>0.13429 | **0.0536**<br>0.04786 |
|  | C | **0.0845**<br>0.08086 | **0.2639**<br>0.14868 | - | **0.0394**<br>0.04845 |
|  | G | **0.2639**<br>0.16051 | **0.0751**<br>0.05139 | **0.0694**<br>0.05139 | - |

## DISCUSSION

Mutation bias across all taxa is typically GC->AT biased (Hershberg and Petrov 2010; Hildebrand, et al. 2010; Liu, et al. 2018) and neutral predicted equilibrium frequencies below GC of 20% (as observed here) are not without precedent (see e.g. (Long, et al. 2018)). Broadly the T enrichment at third sites within the genome is then compatible with a large role for mutation bias, possibly mediated by APOBEC (Simmonds 2020). However, we have shown that nucleotide usage, while skewed in the direction imposed by mutation bias, is nonetheless deviant from it. The difference between observed and expected T3 and TT (Fig 2) proportions are noteworthy. At four fold degenerate sites while C and G usage are close to equilibrium, A is far above and T is far below (T4=50.8%, T4*=64.6%; A4 = 28.95%, A4* = 18.93; C4 = 13.70%, C4* =12.28; G4 = 6.50%, G4* = 4.19%.). We propose that a parsimonious explanation is that the sizeable mutation bias towards T generates deleterious mutations, even at synonymous sites, and selection therefore favours reduced T content. However, increasing C or G potentially comes at a cost of increased CpG, so the base most in excess of its equilibrium is A. As a consequence, while CpG avoidance is real in some genes, GC3 is a little higher than predicted from the underlying mutational profile. This thus presents an unusual case in which the most common synonymous codons (those ending in T) are not the selectively advantageous ones.

There are, however, at least four problems with our mode of analysis. First, a theoretical alternative explanation for the difference between predicted and observed values is that the virus was at neutral mutational equilibrium in its prior host (cf. H1N1, Ebola), but since the transfer to humans the mutational profile has altered. Were this so we may just have identified a lag in viral evolution from one neutral equilibrium to another. In this context deviation from equilibrium has little if anything to say about either selection or optimal vaccine design. While evidence for GC->AT biased mutation in related viruses (Simmonds 2020) renders this less parsimonious an explanation, direct examination of mutational profiles of the virus in its ancestral host (whatever that may be) would be valuable. The evidence for subtly but significantly different mutational matrixes dependent on the class of site employed provides more direct evidence for contemporary selection on T content throughout gene bodies that cannot be accounted for by a temporal shift in mutational profile.

Second, assuming no change to the mutational matrix, *sensu stricto*, we have observed a fixation bias (Lercher, et al. 2002). Evidence for a fixation bias need not necessarily indicate the direction of selection, as selection bias is only one class of fixation bias. In biased gene conversion, for example, the mismatch repair machinery recognises, during double strand break repair, heteroduplex GC:AT mismatches and corrects these in favour of GC residues (Brown and Jiricny 1988). This causes a meiotic drive like process in which deleterious mutations can be driven to higher frequencies (for further consideration see Hurst 2019). Given that single strand RNA cytoplasmic viruses, such as SARS-CoV2, are unlikely to be exposed to the nuclear mismatch repair machinery or need double strand break repair, biased gene conversion is unlikely to explain GC3>GC3*, T3<T3* *etc.*. We cannot with our data, however, rule out unknown mechanisms causing similar non-selective fixation biases. It is then valuable to provide more direct evidence for an advantageous effect of reduced T3/increased GC3, as suggested by our preliminary analysis on expression level. Experimental manipulation of GC3 content (cf. Kudla, et al. 2006; Mordstein, et al. 2020) is a high priority.

Third, we have presumed that the mutational spectrum observed at 4-fold degenerate sites is a good reflection of the true mutational profile. Typically, when applying methodology like this we presume that the temporal proximity between occurrence and observation of mutations is so small that there has been no time for selection to filter in a manner that distorts the mutational matrix. Unusually, however, we found that although slight, there is a difference between the mutational profile observed at CDS sites that are not 4-fold degenerate and those that are. While this difference is so slight it cannot explain why T is so deviant from equilibrium levels, and doesn’t question our overall findings, we do nonetheless presume that the 4-fold site matrix itself is unbiased. If sequencing strains hours apart in time and applying only mutations at 4-fold degenerate sites is biased, this would require so strong selection at 4-fold redundant sites. While not obviously plausible we have no means to disprove this (and strong selection, albeit associated with splicing, has been identified at synonymous sites in human genes (Savisaar and Hurst 2018)).

Fourth, we have presumed that, after filtering (see Methods), all sequences are error free. While sequencing errors cannot explain a bias as strong as the difference between excess and expected TT or T3, nor can they obviously explain the evidence for contemporary selection against T, it may possibly explain the small difference between predicted and observed nucleotide content at 4-fold sites for G and C (the deviations of A and T from predicted equilibria are relatively large). One suggested means to avoid this is to only employ mutations that have been sequenced more than once (Hildebrand, et al. 2010). However, this has been shown to introduce its own bias (Charneski, et al. 2011). Using high quality sequence, it was shown that using mutations that appear once and those that appear twice or more makes a significant difference to the matrix and estimates of equilibria (Charneski, et al. 2011). The cause of this is likely to be a selection filter: mutations that persist longer to be sequenced twice or more will be skewed towards milder effect mutations. This accords with our observation of a slight difference between matrixes that restrict just to 4-fold degenerate sites and those that do not. The ideal then is to filter not be regularity of appearance but by sequencing quality (hence our decisions on which sequences to employ: see Methods). Nonetheless, to err on the side of caution we considered mutations at 4-fold degenerate third sites that appear more than once (i.e. excluding singletons) and found that GC* is now even lower than previously predicted (GC*=10.3%, 95% bounds 10.11 −10.61). Thus, we are confident that we can exclude sequencing error as an explanation for observed GC3 > GC3* and T4<T4* (singleton excluded prediction of T4*=79.5%). Nonetheless, owing to observation bias and low sample size we caution against over-interpretation of this result.

Assuming we have identified the direction of selection (against T, against CpG in some genes) this can inform vaccine design. Unusually, even though T is the most common nucleotide at third sites (by a considerable margin), we propose increasing this even more thereby forcing the viruses against the direction of purifying selection. We predict that raising CpG in the genes that are CpG deficient would be a viable strategy even at a cost of raising GC3/lowering T. By contrast for those few genes with E(CpG)>1 (i.e. gene E, ORF10, see also Digard et al. (2020)) CpG manipulation increasing GC3 would be a dangerous strategy, potentially achieving little more than an increase in expression. Increasing their AT content would appear to be the anti-selection direction. We note however that ORF10’s function, if any remains, unclear there being no evidence of transcripts from it, despite it looking like a well formed ORF (starts ATG stops TAG, multiple of 3 long). Its GC3 content is also far from neutral equilibrium (GC3=36%). In this context gene E may be a good one to alter synonymous site usage as it appears not to be under selection for CpG or TpA avoidance.

Genes ORF1a, ORF1b, ORF6, ORF7b and S are good candidates for the raising of CpG content. Gene N is noteworthy in being very highly expressed, long (1260 bp), GC rich (GC3=38%) and with moderate CpG enrichment (E(CpG)=0.56). Given these characteristics it should be possible to increase CpG by manipulating some third sites (those with C at codon position 2 or G at codon position +1) while reducing GC and increasing T content at other sites. For smaller genes there is less leeway. In this context S, ORF1a and ORF1b are also very strong candidates being long, with moderate GC3 and low CpG enrichment.

## Acknowledgements

This work was supported by the Wellcome Trust (fellowship 207507 to G.K.) and the European Research Council (advanced grant ERC-2014-ADG 669207 to L.D.H.)

## Supplementary files

S Table 1: Ligand binding by antiviral proteins

S Table 2: Data on nucleotide content of viruses

S Table 3 SARS-CoV2 genomes used and acknowledgement

S Table 4 H1N1 genomes used and acknowledgement

S Table 5 H1N1 genomes used and acknowledgement

S Table 6 The dinucleotide mutational matrix for SARS-CoV2

