## Supplementary Table 1 for "Evidence for strong mutation bias towards, and selection against, T/U content in SARS-CoV2: implications for attenuated vaccine design"

|  | **Ligand binding** | **Sequence Binding preference** |
| --- | --- | --- |
| **MDA5** | long dsRNAding activity^1–4^ | May have preference for AU-rich - unclear at the moment ^1,5,6^ |
| **LGP2** | 5' triphoshphate ssRNA, dsRNA^7^ | No known prefernce - binds to termini of ligands^4,7,8^ |
| **RIGI** | 5’triphosphate ssRNA or short dsRNA^3,9^ | poly (U/UC), poly (A/AG) ^10,11^ |
|  |  | AU rich hairpins and short dsRNA ^5,6,12^ |
| **OAS/RNAse L** | dsRNA (OAS)^13^ | OAS1 activated by NNWWNNNNNNNNNWGN motif and GU wobble bases in dsRNA^14,15^  OAS2/3 no known preference |
|  | dsRNA, ssRNA (RNaseL)^16,17^ | Upon activation by OAS, cleaves predominantly after UpU and UpA dinucleotides^17,18^ |
| **PKR** | dsRNA^19^ | No preference^20^ |
| **ILF3** | dsRNA, ssRNA^21–23^ | AREs ^24^ |
| **TRIM25** | dsRNA, ssRNA^25^ | GC rich sequences^25^ |
| **ADAR** | dsRNA^26^ | No sequence binding preference but editing preference depending on ADAR protein^27^ |
| **ZAP** | ssRNA^28,29^ | CpG^30^ |
| **DDX17** | hairpins on ssRNA^31^ | CA and CT repeats^31^ |
| **cGAS** | cytoplasmic dsDNA^32^ | short region of dsDNA flanked with at least 3 G^32^ |
| **DAI** | Cytoplasmic DNA ^33^ | No preference^34^ |
| **IFI16** | Cytoplasmic DNA^35^ | No preference^36^ |

**Cytplasmic PRRs**
